# Is this scenery worth exploring? Insight into the visual encoding of navigating ants

**DOI:** 10.1101/2024.08.28.610048

**Authors:** Leo Clement, Sebastian Schwarz, Blandine Mahot-Castaing, Antoine Wystrach

## Abstract

Solitary foraging insects like desert ants rely heavily on vision for navigation. While ants can learn visual scenes, it is unclear what cues they use to decide if a scene is worth exploring at the first place. To investigate this, we recorded the motor behavior of *Cataglyphis velox* ants navigating in a virtual reality set-up and measured their lateral oscillations in response to various unfamiliar visual scenes under both closed-loop and open-loop conditions. In naturalistic-looking panorama, ants display regular oscillations as observed outdoors, allowing them to efficiently scan the scenery. Manipulations of the virtual environment revealed distinct functions served by dynamic and static cues. Dynamic cues, mainly rotational optic flow, regulated the amplitude of oscillations but not their regularity. Conversely, static cues had little impact on the amplitude but were essential for producing regular oscillations. Regularity of oscillations decreased in scenes with only horizontal, only vertical or no edges but was restored in scenes with both edge types together. The actual number of edges, the visual pattern heterogeneity across azimuths, the light intensity or the relative elevation of brighter regions did not affect oscillations. We conclude that ants use a simple but functional heuristic to determine if the visual world is worth exploring, relying on the presence of at least two different edge orientations in the scene.

**Summary statement:** Using a virtual reality setup, we reveal that ants rely on a heuristic to trigger visual exploration in an unfamiliar scene. The simultaneous presence of vertical and horizontal edges is necessary and sufficient for the ants to produce lateral oscillations and scan the scene.

## Introduction

How can insect navigate complex natural environments using vision despite their small brain and low-resolution eyes has been the focus of decades in research (Baddeley et al., 2011, 2012; Collett et al., 2013, 2007; Collett and Cartwright, 1983; Cormons and Zeil, 2023; Franzke et al., 2020; Graham and Cheng, 2009; Graham and Collett, 2002; Hoinville and Wehner, 2018; Kohler and Wehner, 2005; Konnerth et al., 2023; Mangan and Webb, 2012; Menzel et al., 2005, 2018; Moura et al., 2023; Philippides et al., 2011, 2013; Thomson, 1996; Wehner and Räber, 1979; Woodgate et al., 2016b; Wystrach et al., 2011a, 2013; Zeil et al., 2003; Zeil, 2012). We know that the visual systems of insects extract specific features such as boundaries and edges (Harris et al., 2007; Horridge, 2009; O’Carroll, 1993; Seelig and Jayaraman, 2013) as well as relative brightness and colour information (Ernst and Heisenberg, 1999; Horridge, 2005; Von Frish, 1914). Although insects can recognise specific, ecologically pertinent objects with which they interact, such as flowers for foraging bees (Chittka and Raine, 2006) or competitors for territorial hover flies (Nordström and O’Carroll, 2006), evidence shows that when it comes to visual navigation, the perceived scene is recognised as a whole through a wide and low-resolution input spanning their whole panoramic views, which naturally encompass the local landmarks and distal panorama without the need to extract them as individual objects (Buehlmann et al., 2016; Graham and Cheng, 2009; Graham and Philippides, 2017; Stürzl et al., 2016, 2008; Towne and Moscrip, 2008; Wystrach et al., 2011a, 2011b; Zeil, 2012; Zeil et al., 2003). Notably, navigating insects rely on the overall shape of the panoramic skyline – the border between terrestrial objects and the sky (Collett et al., 2007; Fukushi, 2001; Graham and Cheng, 2009; Reid et al., 2011) or the position of large dark areas on their retina rather than their specific, local shapes (Buehlmann et al., 2016; Ernst and Heisenberg, 1999; Horridge, 2005; Lent et al., 2013a; Woodgate et al., 2016a). In addition to these static cues, bees, wasps and to a lesser extent ants can also use dynamic cues like translational optic flow for guidance (Dittmar et al., 2010; Esch et al., 2001; Ronacher et al., 2000; Ronacher and Wehner, 1995; Srinivasan et al., 2000, 1996; Stürzl et al., 2008).

To characterise the visual cues used by navigating insects, some studies exploited the ability of experienced foragers to use learnt visual information (Chaib et al., 2021; Cormons and Zeil, 2023; Dittmar et al., 2010; Horridge, 2005; Jayatilaka et al., 2013; Kohler and Wehner, 2005; Konnerth et al., 2023; Mangan and Webb, 2012; Moura et al., 2023; Murray et al., 2020; Narendra et al., 2013; Philippides et al., 2013; Towne et al., 2017; Wystrach et al., 2013), while other studies focused instead on the spontaneous orientation response of naïve individuals in relation to particular visual features (Bausenwein et al., 1994; Buehlmann et al., 2020b; Buehlmann and Graham, 2022; Byrne et al., 2003; el Jundi et al., 2016; Franzke et al., 2020, 2022; Goulard et al., 2021; Grabowska et al., 2018; Graham et al., 2003; Heusser and Wehner, 2002; Maimon et al., 2008; Philippides et al., 2013; Wallace, 1958, 1959; Woodgate et al., 2016b). In both cases, these approaches revealed the nature of the cues used for heading control, that is, when the insect uses visual cues to maintain its course towards a particular direction, learnt or not (Buehlmann et al., 2020a; Grabowska et al., 2018; Graham et al., 2003; Harris et al., 2007; Heusser and Wehner, 2002; Judd and Collett, 1998; Pratt et al., 2001; Stürzl et al., 2016; Wallace, 1959; Zeil, 1993).

Here, we did not investigate which features are important for heading control. Instead, we focused on whether some features need to be present within a visual scene, unfamiliar to the ants, to trigger an exploratory behavior in the first place. In other words, does an exploratory behavior occur ‘by default’, that is, as soon as the insect is outside of its nest, even if it is still in the dark, or does it require the presence of specific visual features? Are such features simple and general such as the mere presence of light, horizontal or vertical edges or does their diversity and position also matter? Would an insect explore an artificial pattern repeated across the whole 360° scene in the same way as a natural-looking scene containing an heterogenous diversity of information across azimuths? Does the insect need to perceive the presence of some general signatures of natural scenes, such as a darker ground and a brighter sky? Overall, do insects use a visual filter to determine whether the surrounding scenery is worth attending to for exploration? We know that insects encode celestial cues through a ‘matched filter’ (Cheng and Freas, 2015; Wehner, 1987), that is, a neural filtering that ‘expects’ and echoes aspects of the natural spatial organization of the polarized light pattern across the sky (Aepli et al., 1985; Homberg, 2004; Homberg et al., 2011; Zittrell et al., 2020). Here, we wonder whether navigating ants possess an equivalent ‘matched filter’(Cheng and Freas, 2015; Wehner, 1987), which would filter natural aspects of terrestrial scenes and hence serves to only trigger the motivation to explore when appropriate.

We used a virtual reality (VR) setup to record the detailed motor behaviour of the solitary foraging ant, *Cataglyphis velox,* when presented with various visual sceneries (video S1). We quantified their lateral oscillations, which many insects spontaneously display when navigating (Cheng, 2022; Clement et al., 2023; Dauzere-Peres and Wystrach, 2023; Freas and Cheng, 2022; Iwano et al., 2010; Kanzaki et al., 1992; Kuenen and Baker, 1983; Namiki and Kanzaki, 2016a; Olberg, 1983; Wystrach et al., 2016). In ants, these oscillations are crucial for visual navigation (Clement et al., 2023; Lent et al., 2013b, 2010; Murray et al., 2020). Oscillations are tuned to enable the ants to scan multiple directions while walking and are upregulated when in an unfamiliar environment (Clement et al., 2023; Murray et al., 2020) or when exploring a novel scene during naive ants’ early learning walks (Clement et al., 2023; Jayatilaka et al., 2018; Zeil and Fleischmann, 2019). Therefore, we used these oscillations as a proxy to measure the ants’ motivation to explore new visual surroundings. This approach allowed us to investigate whether the exploratory motivation depends on the visual structure of the environment and to identify which visual cues prompt exploration.

Our results highlight the importance of specific static visual features, notably the simultaneous presence of a diversity of edge orientations, to trigger the production of regular oscillations. Dynamic features, such as rotational optic flow, turn out to be important for the oscillation’s amplitude control but not for their production. These findings are discussed in the context of the established neural circuitry and visual ecology in ants.

## Materials and Methods

### Study Animal

We used the Iberian thermophilic desert ant species *Cataglyphis velox.* In this species, foragers do not use pheromone tracks but forage solitarily (Cerdá and Retana, 2000) relying mainly on learnt terrestrial visual cues and path integration (PI)(Mangan and Webb, 2012).

Three nests of the species *C. velox,* were collected in Seville (Spain). Since 2020 those nests were maintained at the lab at the Paul Sabatier University, Toulouse. The ant colonies were housed in vertical nests created by excavating galleries and chambers in aerated concrete. These nests were maintained with controlled ventilation, temperature (24-30°C), and humidity (15-40%). Each nest was connected by a 20 cm transparent tube to a 40 x 30 cm foraging arena with sand on the floor and without any specific visual enrichment beyond the water tubes within the arena and the view of the experimental room beyond the arena’s 10 cm high walls. The foraging arena was exposed to a heating lamp, a natural 12-hour day-night light cycle that included ultraviolet light (UV). During data collection, colonies were underfed to ensure their motivation to search for food.

### Trackball set-up & Virtual Reality

To record the ants’ movement, we mounted them on a trackball device (Dahmen et al., 2017). This device consists of a polystyrene ball held in levitation in an aluminium cup by a stable air flow. The trackball has two sensors placed at 90° to the azimuth of the sphere, which record the movements of this sphere and translate them into X and Y coordinates retracing the path of the ant (Fig. 1A). The X and Y acquisition of the trackball rotations happened at a 30 fps, enabling us to reconstruct the ant’s movements with high precision. Ants were tethered and placed on top of the ball. The tether consisted of a 0.5 mm pin that could rotate within a vertical glass capillary positioned above the ant (as in Dahmen et al., 2017) and attached to the ant thorax, to which we applied a drop of magnetic paint, via a micro-magnet. This tether allows the ants to rotate their bodies in the yaw axis while maintaining them on the top of the trackball (Fig. 1A). The polystyrene ball was prevented from rotating in the horizontal plane by two small vertical wheels touching the ball’s equator. As a result, ants could physically control their heading direction on top of the ball, but translational movements resulted in rotations of the ball. This mounting device provides a more natural experience for the ants. First, it has the advantage of naturally coupling body rotations with the expected visual feedback, without the computing lags of needing to rotate the visual scene around the animal. Second, the force the ant must produce to rotate their body on top of the ball is equivalent to when walking on the floor. By comparison, more traditional tethers where insects are fixed require around a hundred-fold increase in force to rotate the ball along the horizontal plane (Dahmen et al., 2017). Finally, ants, mounted on the same trackball set-up in this ‘free to rotate’ tether, are known to use learnt terrestrial cues, perform path integration and search when in unfamiliar environments in a remarkably similar manner as when they are on natural grounds (Clément et al., 2016). Also, the amplitudes and frequencies of oscillations recorded here corresponds to what is observed in the field with ants of the same species running on natural ground (Haalck et al., 2023). Together, this provides conclusive evidence that our trackball setup does not disturb or prevent the ant’s natural navigational behavior.

**Figure 1:**
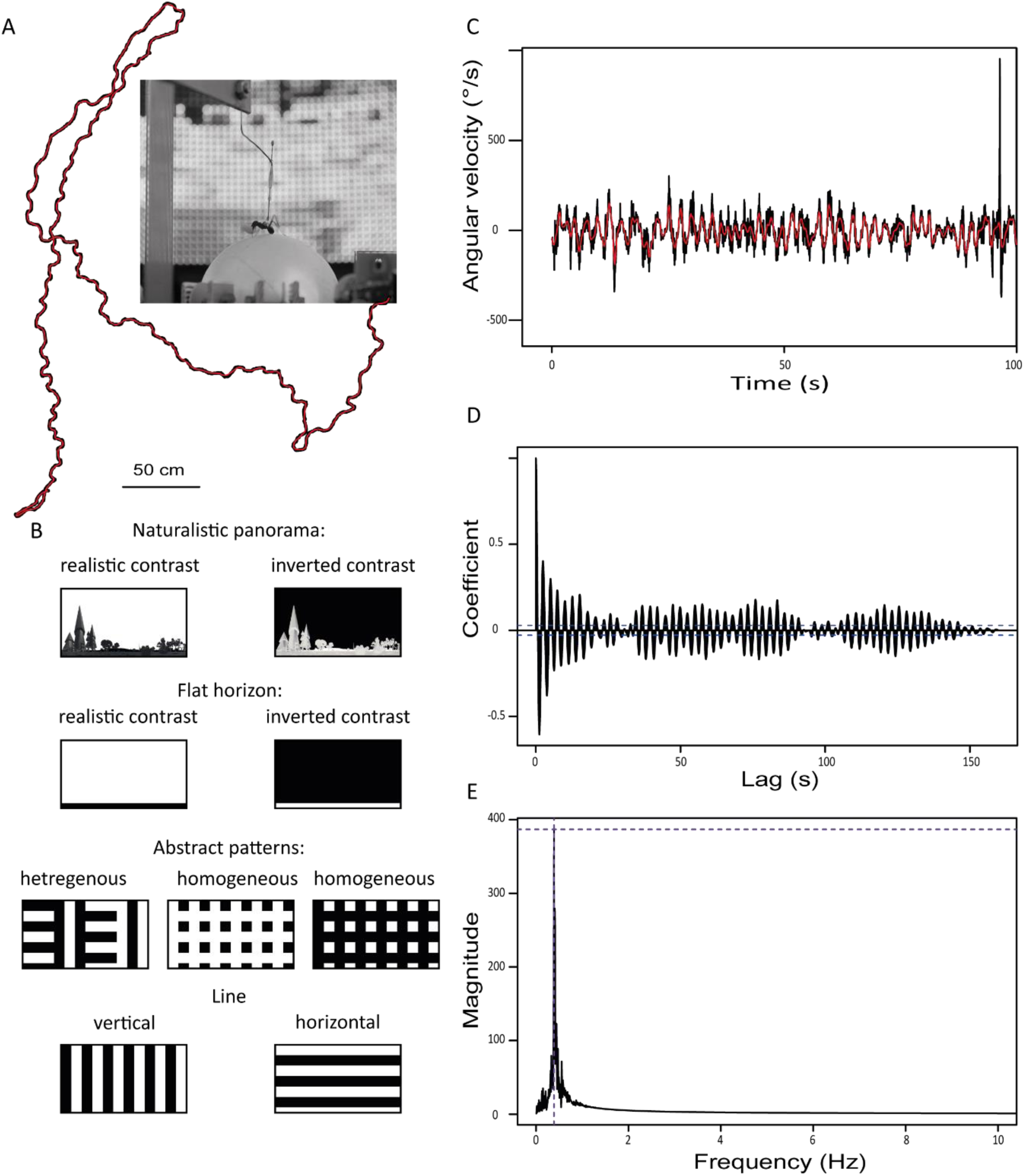
Trackball set-ups, recording and processing of the ants’ trajectory. (A) Picture of the trackball within the virtual reality set-up (side view). Two wheels prevent the sphere from rotating along the horizontal plane. Ants are free to rotate their body along the yaw axis to control in which direction they perceive the world. The path shows an example of ants recorded in a naturalistic environment (from B). Black path is the original recording, the overlay red path is the smoothed one (Savitzky-Golay filter of 2 s) (B) Different background displays (conditions) during the experiment. (C) Angular velocity signal over time of an individual example path (from A) with smoothed signal superimposed (red). (D) Autocorrelation carried out on the entire smoothed angular velocity signal. (E) Fourier transformation of the autocorrelation coefficients signal (shown in D) provides the ‘power spectral density’. This approach has the advantage to provide magnitudes that are directly comparable between individuals. For each individual, the frequency peak with the highest magnitude was extracted, indicating a strong oscillation of the angular velocity signal at that frequency (dashed blue lines).

During experiments, ants on the trackball were immersed at the centre of a 360° visual display (Fig. 1A): a cylinder of 50 cm diameter and 76 cm high, which inner surface is covered by LED panels (73728 LEDs) providing a light intensity up to up to 860 mcd per pixel and spatial resolution of 0.94 degrees per pixel, which exceeds the visual acuity of *Cataglyphis* eyes (>2 degrees per pixel (Zollikofer et al., 1995)). The LEDs are controlled through the computer running a virtual environment using the freeware Unity 2020.1.

In this study, we used two different type of conditions hereafter referred to as ‘open-loop’ and ‘closed-loop’ experiments, respectively.

#### ‘Closed-loop’

The translational displacement of the ants modifies the VR display, by updating its position in the 3D reconstructed world according to its displacement. This generates translational optic flow through the floor texture and the nearby objects’ apparent movement. Note that since the ants can rotate on top of the ball, it is not needed to rotate the scene, and the point of expansion of translational optic flow is directly linked to the direction of rotation of the ball, that is, to the ants’ actual movement direction.

#### ‘Open-loop’

The ants’ movements do not modify the virtual reality display, which thus displays only a fixed image. As a result, there is no translational optic flow in this condition, however the rotation of the ants on top of the ball will naturally produce the expected rotational optic flow on their retinas.

### Virtual environment & experiments

To identify the key visual features that trigger exploration, we recorded ants in various unfamiliar environments, spanning from natural sceneries to abstract patterns (Fig. 1B). The natural looking panorama consisted of an infinitely distant horizon displaying a skyline recorded in a typical natural habitat of these ants in Seville, as well as a hundred or so individual objects (trees, bushes, rocks) spread over an area corresponding to one hectare (100 x 100 m) in relation to the ants’ physical displacement. The ratios between the object sizes and the ant’s movements were similar to natural conditions, with the ant’s viewpoint placed at 5 mm above the ground, and the biggest trees extending up to 30 m in height. The ants started the assay at the centre of this world, which was thus large enough to ensure that, when in closed-loop condition, they could not reach the world’s borders within the experiment’s duration (160 sec). Importantly, these ants had no prior experience of the VR setup, nor of the world displayed. Each ant was tested once in up to three different visual conditions in a random order. Between each test, ants were released inside their nest for at least 3 min before being retrieved for another trial in a different visual condition. Each individual was therefore placed in the VR up to three times but was only used once for each visual condition.

### Data extraction and analysis

All statistical analyses were run using the free software R (v 3.6.2. R Core Development Team).

To determine the presence of regular lateral oscillations, our focus was on the angular velocity, which is a direct measurement of the left/right motor control. We processed the time series data through three successive steps to calculate its spectral density using the Wiener-Khinchin theorem. First, we smoothed the X, Y path (Fig. 1A, red path) using a Savitzky-Golay filter (from R “trajr” package) with a window length of 2 s. Then, we extracted the angular velocity time series, which was smoothed twice: first with a moving median, then with a moving mean, both with a window length of 0.5 s (Fig. 1C). Next, we conducted an autocorrelation function (Fig. 1D) on the smoothed time series and performed a Fourier transformation on the autocorrelation coefficient to obtain the power spectral density (Fig. 1E). For each individual, we extracted the dominant frequency (i.e., the frequency with the highest peak magnitude) and the corresponding magnitude (Fig. 1E). To determine if these magnitudes indicated a significant regular oscillation, we compared them to an ‘average’ spectral density magnitude obtained by resampling randomly the ant angular velocity time series a hundred times. These resampled signals underwent the same processing as the original angular velocity time series data, including smoothing, autocorrelation, Fourier transformation and extraction of the highest peak magnitude. For each ant, the 100 highest magnitudes obtained were averaged and then compared to the individual’s real signal magnitude using a Wilcoxon one-tail test for paired data.

### Statistical models

To determine which key features has an impact on the oscillation behaviour, we compared our conditions using mixed models. Two types of models were used: one that considers the interaction and the other one with simple additive effects. If the residuals of these models deviated from normality and/or homoscedasticity, the response variables were transformed. The model was selected and further analysed through an analysis of variance (Anova), followed by a Tukey’s rank comparison post-hoc analysis. Additionally, multiple independent experiments were performed and each individual was tested across different conditions. As a result, the models were mixed models that controlled for the effects of the sequence and individual, as well as considering the set of experiments as a random variable.

## Results

We recorded the motor behavior of navigating *C. velox* ants, captured in their foraging area next to their nest. Crucially, these ants had no prior experience with the VR setup and were tested in different unfamiliar visual conditions only once to prevent the influence of familiarity visual recognition signals. Each ant was individually placed into the VR by tethering them on top of the trackball in a way that enable them to physically rotate and control their actual body orientation as when walking on the ground, as in (Clement et al., 2023; Dahmen et al., 2017; Murray et al., 2020). The resulting paths obtained showed regular lateral oscillations – that is, an alternation between left and right turns at a steady rhythm along their path – visible by naked eyes (Fig. 1A). To quantify the regularity of these oscillations we extracted the angular velocity signals form the paths and computed the Fourier’s power spectral density (PSD; Figs. 1, S1). The magnitude of the PSD Fourier’s peak, which represented the most prominent rhythm in the signal, reflects the regularity of the oscillations for a given frequency, with higher magnitudes indicating a more consistent rhythm.

### Ants do oscillate in the virtual reality with a naturalistic reconstructed environment

We first investigated whether ants would display oscillations in the VR when in closed-loop with a realistic-looking unfamiliar virtual environment. The 3D world consisted of a reconstruction of a typical panorama of the ant’s habitat (Fig. 1B, first picture) providing a rich distant panorama (horizon line) as well as distant and proximal cues (bushes, trees, rocks, etc.). In this closed-loop condition the ant experienced translational and rotational optic-flow in response to its own movement velocities (up to 544 deg/sec for rotations and 26 cm/sec for translation) similarly to what they would experience in a natural scenario. Ants recorded in this reconstructed unfamiliar environment displayed path oscillations visible to the naked eye (Fig. 2A). The highest PSD magnitudes obtained from the ant’s signal were greater than those obtained from randomly resampled signals (Wilcoxon one-tail test: V=15, *P*<0.001; mean±se: naturalistic panorama=179.938±13.53; resampled signal=108.102±0.494), indicating that ants displayed lateral oscillations with a significantly higher regularity than would be expected by chance.

**Figure 2.**
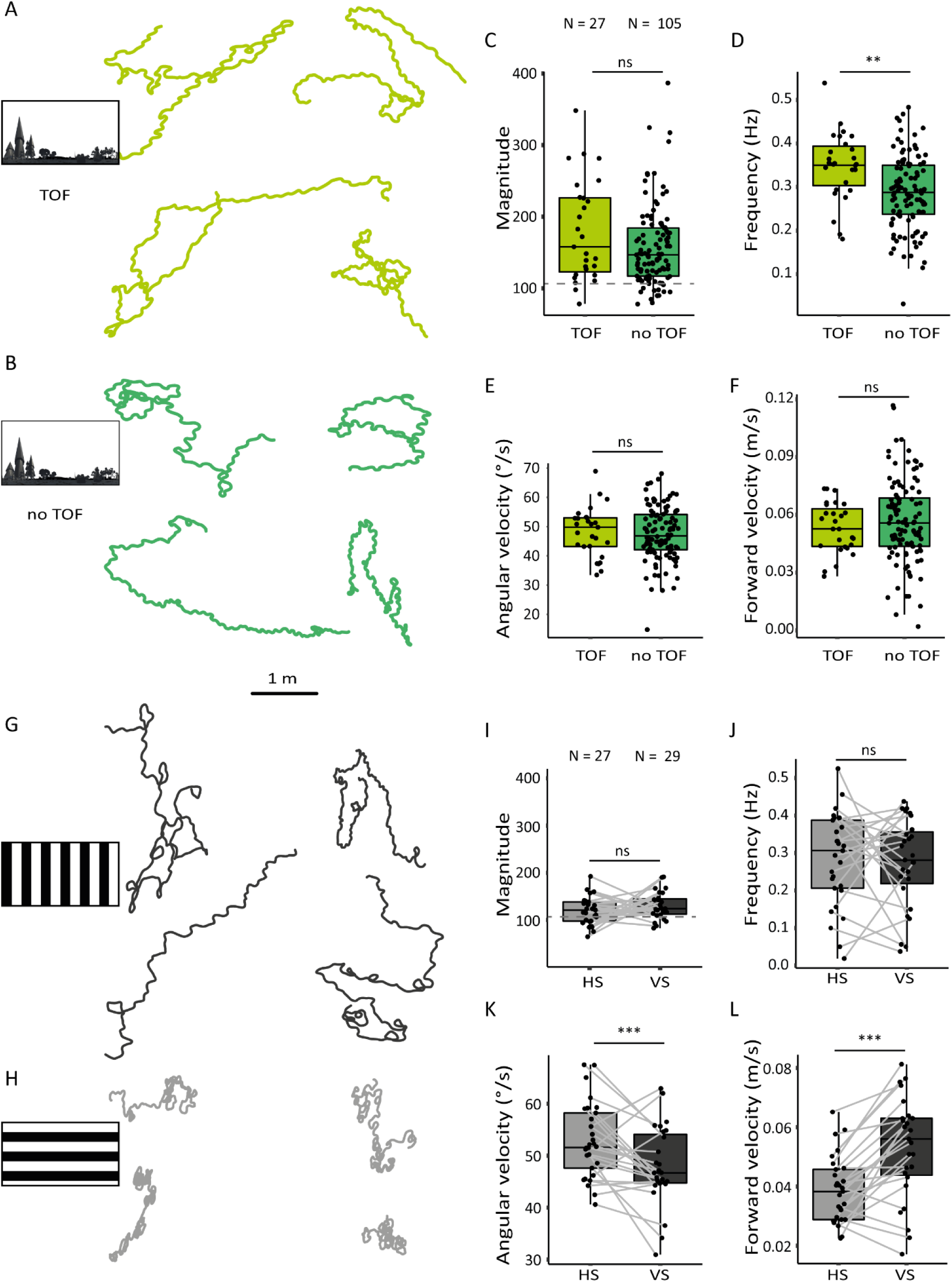
Dynamic visual cues are involved in the control of oscillations. (A-B, G-H) Examples paths of four ants across 160 s recorded with translational optic flow (TOF; A, light green) and without TOF (B, green) or recorded in visual surrounding made of vertical (G, grey) and horizontal (B, light grey) stripes. (C-D; I-J) Distribution of the individual Fourier dominant peak obtained from angular velocities times series across 160 s. High magnitudes indicate a strong presence of this oscillation. High frequencies indicate a fast-oscillatory rhythm. The dashed black line represents the mean of the spectral density peak magnitudes resulting from 100 random resamples of the angular velocity time series (see methods). (C; I) Dominant frequency. (D; J) Highest peak magnitude. (E-F, K-L) Distribution of the angular (E; K) and forward velocities (F; L).

The frequency peak fell within the expected range of 0.1 to 0.54 Hz (Fig. 2D, J), consistent with previous observations in other insect species (Clement et al., 2023; Lonnendonker, 1991; Wystrach et al., 2016). Importantly, this rhythm is 10 to 50 times slower than the ants’ typical stepping frequency (Zollikofer, 1994), showing that it is not a by-product of their walking gait but the result of an independent internal oscillatory mechanism (Clement et al., 2023). Overall, these results confirm that ants produce regular lateral oscillations while walking within this naturalistic-looking virtual environment.

### Dynamic visual cues are involved in the control but not the production of oscillations

In the previous experiment, ants were in closed-loop with the environment, that is, the ant’s movements on the trackball generated translational and rotational optic flow responses in the virtual scene, as well as parallax movements of the proximal objects.

To expand upon this finding, we investigated whether these optic flow cues are necessary to trigger regular oscillations by recordings ants with a static image of the same realistic looking virtual environment (Fig. 2B for example path). Note that by doing so, the ants still perceived rotational optic flow caused by their own rotation on the ball but no longer experienced translational optic flow or parallax movements in response to their movement. In this situation, ants still displayed clear oscillations (Fig. 2B, E; Wilcoxon one-tail test: V=305, *P*<0.001, mean+se: no TOF=158±4; resampled signal=107±0.2). Angular and forward velocities as well as the magnitude (i.e., regularity) of the oscillations were similar to the closed-loop conditions (Fig. 2C, E, F; Magnitude (Fig. S1A, B): F_1,132_=2.2464, *P*=0.134; Angular velocities: F_1,132_= 0.0176, *P*=0.895; Forward velocities: F_1,132_= 0.3852, *P*=0.535), however, the oscillations’ frequency was significantly lower in open-loop condition (Fig. 2D; F_1,132_= 10.801, *P*<0.01; Fig. S1A, B). This shows that translational optic flow does not influence the regularity or production of oscillations but impacts their frequency.

We next tested the impact of rotational optic-flow (ROF) on oscillatory signatures by comparing the ants’ behaviour within abstract visual sceneries consisting of either vertical or horizontal stripes (Fig. 1B, last row), with the latter condition producing no ROF as the ant rotates on the ball. In these abstract environments, ants still display regular oscillations visible to the naked eye and significantly above those that would be expected by chance (Fig. 2G, H; Wilcoxon one-tail test=*Ps*<0.03; means±se: horizontal: 119.535±5.37; vertical: 130.504±5.54; resampled signal both group=107.505±3.89). The absence of rotational optic flow (i.e., with horizontal stripes) dramatically increased the angular velocity (Fig. 2K; F_1,56_= 10.54, *P*<0.001, mean±se: horizontal: 52.776±1.323 °/s; vertical: 47.876± 1.433 °/s) and reduced the forward speed (Fig. 2L; F_1,56_= 26.74, *P* <0.001, means±se: horizontal: 3.86± 2.312 cm/s; vertical 5.285 ±2.504 cm/s), leading the ants to often execute full loops (Fig. 2H). This confirms that ROF is involved in limiting the amplitude of the oscillations, as previously suggested (Clement et al., 2023; Dauzere-Peres and Wystrach, 2023). However, the oscillations’ magnitude (i.e., regularity) and frequency were not different between the vertical or horizontal stripes conditions (Fig. 2I, J; Anova, magnitude: F_1,56_=2.01, *P*=0.155; frequency: F_1,56_= 0.139, *P*=0.708; Fig. S1, C, D), showing that the presence or absence of ROF does not impact the production of oscillations.

### Diversity of edge orientation impacts the production of oscillations

Despite being tested within an abstract environment (presented only vertical or only horizontal edges), ants still displayed clear oscillations. However, these were less regular (lower PSD magnitude) than those produced in the naturalistic-looking environment (naturalistic with TOF=180±12; naturalistic no TOF=158±4; horizontal: 119.535±5.37; vertical: 130.504±5.54). We first hypothesized that this was due to the lack of heterogeneity across heading directions of these abstract visual scenes presenting a regular pattern of stripes, which would decrease the motivation to explore and inhibit the production of oscillations. Indeed, functionally, a visual environment consisting of a homogeneous pattern across azimuths is less worth exploring than a heterogeneous one. To test for this, we recorded ants in visual sceneries presenting a same number and diversity of edges (both vertical and horizontal), but with either a homogeneous or heterogeneous distribution of these stripes across azimuths (Fig. 3A). We found no effect on either the regularity or frequency of the oscillations across these different experimental conditions (F_1,59_= 0.1253, *P*=0.722), showing that the homogeneity/heterogeneity of the visual scene across azimuth had no impact on the production or control of oscillations.

**Figure 3.**
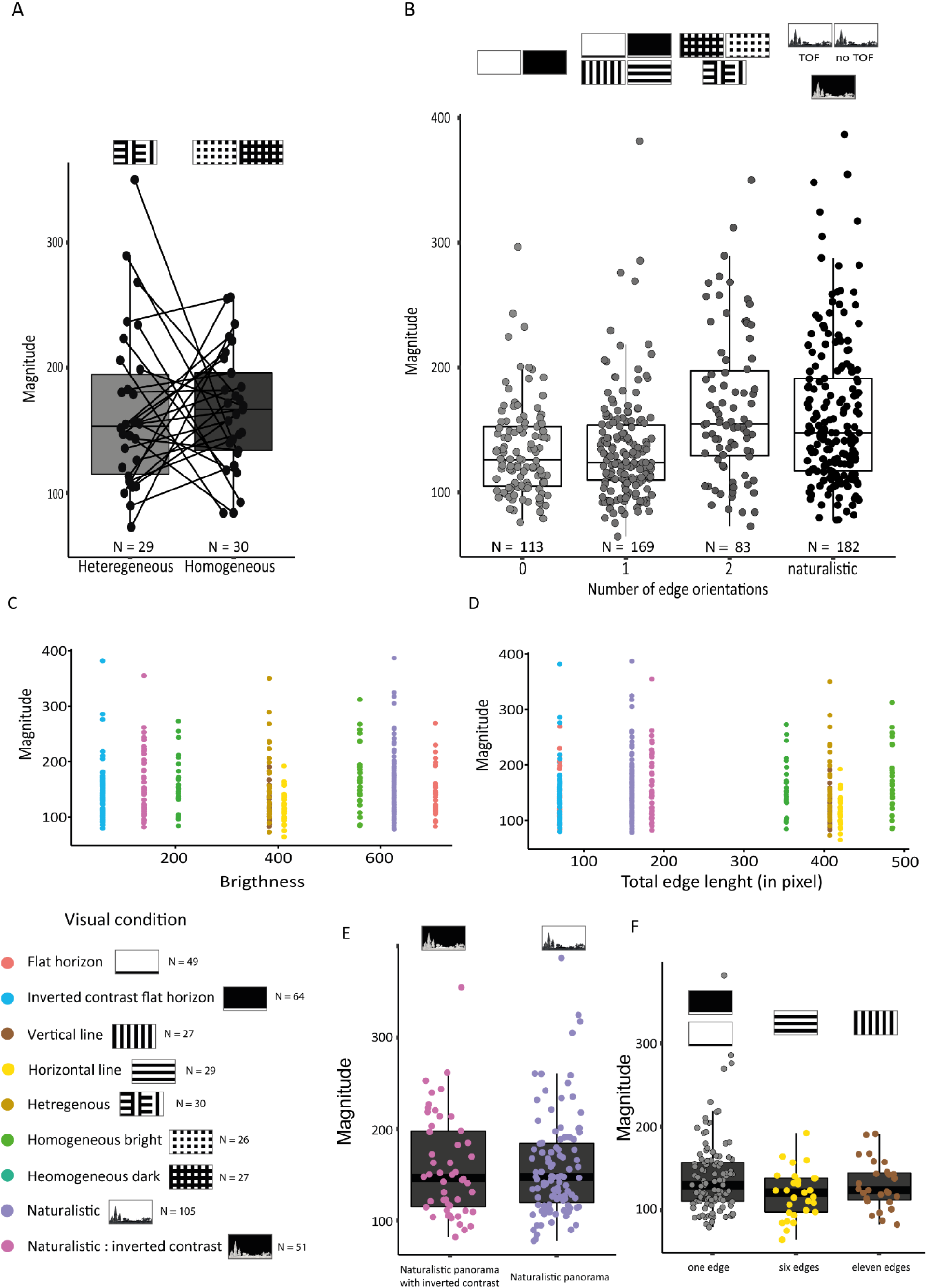
Ants display regular oscillations in visually structured environments. (A, B) Distribution of the individual Fourier dominant magnitudes obtained on angular velocities times series across 160 s. High magnitudes indicate a strong presence of this oscillation. (A) Ants recorded in heterogeneous (left) or homogeneous (right) background constitute of horizontal and vertical stripes. (B) Pooling of several experimental conditions according to the number of edge orientations from absence of edges to several. **(C-F)** Distribution of magnitudes according to (C) the brightness, (D) length of edges, (E) position of contrast and (F) number of stripes. For (C-F) the diversity of edges orientation significantly impacts the oscillations magnitude whereas the brightness (C) or the total edge length (D) did not have an effect (Anova: edge orientation *Ps*<0.05, other parameters *Ps*>0.05).

However, the magnitude (i.e., regularity) of oscillations in these still artificial conditions was similar to the one recorded in a naturalistic visual scenery (mean±se: naturalistic: TOF=180±12; no TOF=158±4; heterogeneous=164±12; homogeneous: 165±9), suggesting that the two different edge orientations (vertical *and* horizontal) are sufficient for the ants to fully produce oscillatory behaviours. Based on this, we put forth the hypothesis that a simultaneous presence of diversity of edge orientations plays a role in the production of oscillations.

To test this idea, we compiled and analysed the results of our conducted series of experiments with a variety of visual backgrounds, spanning from visual scenes, which contained a complete absence of edges (uniform black or uniform white), scenes containing a single edge or two edge orientations (both horizontal and vertical) or complex visual scenes with many edges (Fig 3B). Across these four conditions, the number of edge orientations had a strong impact on the regularity of oscillations (F_3,547_= 40.559, *P*<0.001, mean±se: zero=133.22±3.483; one=134.519±3.223; two=167.4067±6.4; naturalistic: 161.4075±4.326; Fig. 3,). Contrastingly, the overall brightness (Anova: brightness: *χ*^2^ = 0.775, P=0.379; number of edges orientation: *χ*^2^=19.050, *Ps*<0.001; Fig. 3C), the actual total edge length across the scenery (Anova: edges total length: *χ*^2^=1.62, *P*=0.203; number of edge orientations *χ*^2^=25.925, *Ps*<0.001; Fig. 3E) and the position of contrast (Fig. 3E, F; see inverted contrast condition) of the pattern had all no significant effect on the observed oscillatory characteristics.

Thus, for ants, the primary visual features that trigger the production of regular oscillations appears to be based on the simultaneous presence of a diversity of edge orientations rather than on the other parameters we tested (Fig. 3 C-F).

## Discussion

We demonstrated that ants navigating in a VR environment exhibit their natural oscillatory behaviour, with magnitudes and frequencies that closely resemble those observed in ants navigating naturally in their environment (Clement et al., 2023; Jayatilaka et al., 2018; Murray et al., 2020). Such oscillations reflect the activity of an intrinsic neural oscillator, widespread across species and serving various navigational contexts (Clement et al., 2023; Dauzere-Peres and Wystrach, 2023; Iwano et al., 2010; Lonnendonker, 1991; Namiki and Kanzaki, 2016a; Steinbeck et al., 2020; Wystrach et al., 2016). These oscillations are key to visual navigation as they optimize visual exploration and are amplified when the ant is in an unfamiliar visual environment (Clement et al., 2023). Therefore, we reasoned that the production of such oscillations can serve as a proxy to quantify the ant’s motivation at exploring the visual world. This provided us with a tool to investigate whether the exploratory motivation in ants depends on the visual structure of the surroundings. Correspondingly, we identified which visual cues are key for the foraging ant to trigger such an exploratory behaviour. We used VR sceneries, unfamiliar to the ants, to prevent the influence of familiarity visual recognition signals on left and right turns (Wystrach et al., 2020) and thus observe a more direct expression of the oscillatory behaviour (Clement et al., 2023). However, our conclusions regarding the up- or down regulation of the oscillator, based on the structure of the scene and its control by dynamic cues, may well apply to both unfamiliar and familiar environments.

### The role of dynamic cues for the control of oscillations

The lateral oscillations and forward displacements of ants in the world generate a dynamic, re-afferent sensory feedback in the form of rotational and translational optic flow, respectively. Such dynamic cues are known to play a major role in insects for various tasks such as speed control (Baird et al., 2010, 2005; Barron and Srinivasan, 2006; Portelli et al., 2011), optomotor responses (Krapp, 2000; Nityananda et al., 2017), distance estimation for landing (Lehrer et al., 1988; Ruffier et al., 2019; Srinivasan et al., 2000, 1996, 1989) the recognition of a familiar scenery itself (Dittmar et al., 2010; Zeil, 1993) or to from a functional compass representation (Beetz et al., 2022). As expected, manipulating these dynamic cues in our virtual reality set-up influenced the dynamics of the oscillations. As previously shown (Busch et al., 2018; Dauzere-Peres and Wystrach, 2023; Egelhaaf, 2023; Franzke et al., 2022, 2020; Pansopha et al., 2014), the absence of rotational optic flow led to a dramatic increase of the turn’s amplitude, up to non-adaptive full loops (Fig. 2H, K) confirming the role of rotational optic flow to control the extent of the current turn. Removing translational optic flow and the parallax motion of the objects – generated by forward motion – down-regulated the oscillation frequency (Fig. 2D). However, neither of these dynamical cues impacted the oscillation regularity (Fig. 2G-I). Overall, this suggests that dynamic cues are dedicated to the control of the dynamics (amplitude and timing) of oscillations but not involved in their actual production.

### The role of static cues for the triggering of oscillations?

Interestingly, presenting artificially impoverished visual sceneries disrupted the regularity of the oscillatory rhythm of the ants (Fig. 3B), showing that the structure of the world perceived by the ants influences the production of oscillations. The crucial parameter to explain the presence of prominent, regular oscillations turned out to be the diversity of edge orientations (Fig. 3B) rather than the overall brightness, the distribution of the contrasts, the quantity of edges in the scene (Fig. S2) or the heterogeneity of the visual pattern across the azimuths (Fig. 3A). For instance, oscillations are equally downregulated in a uniform background (black or white Fig. 3B), slightly more prominent in a scenery consisting of only one type of edge orientation (horizontal or vertical and independently of the number of bars displayed) and fully reinstated in sceneries with a combination of at least two edge orientations (Fig. 3B). The insect’s visual system is known to extract edges to detect the boundaries of objects and surfaces (Buehlmann and Graham, 2022; Grabowska et al., 2018; Horridge, 2009; Maimon et al., 2008). Here we show that the presence of this feature, at least in *C. velox* ants, is important to trigger oscillations and thus a proper exploration of the scene.

Ants are well-known to use the skyline to maintain their direction of travel (Collett et al., 2007; Fukushi, 2001; Graham and Cheng, 2009; Philippides et al., 2011). The skyline represents the boundary between the sky, on top and the terrestrial objects, below. Given that in natural conditions the sky is generally brighter than the ground, we could have expected that insect possess a ‘matched filter’ (Cheng and Freas, 2015; Wehner, 1987) to process preferentially sceneries that are brighter in the upper part (Möller, 2002; Philippides et al., 2011; Schultheiss et al., 2016). Perhaps surprisingly, inversions of the scenery contrast did not influence the behavioural outcome as ants displayed equally prominent oscillations whether the upper part of the scenery was bright or dark (Fig. S2C). However, the sceneries presented here were monochromatic (black and white) using LED with RGB wavelength (peaks at 624, 520 and 470 nm respectively), and thus lacking UV- as well as polarized light, which is present in natural skylines. Arthropod eyes (Aepli et al., 1985; Labhart and Meyer, 2002; Mote and Wehner, 1980) are sensitive to UV- and polarized light (notably UV polarization) with the latter being detected by ommatidia pointing towards the sky (Aepli et al., 1985; Labhart and Meyer, 2002, 1999; Möller, 2002) and both celestial cues are important for navigational behaviours. Hence, whether ants possess a matched filter for sceneries with UV and/or polarized light in the upper part of their visual field to regulate their oscillations remains to be explored. However, we show here that the direction of edges in the RGB range, which is picked up mainly by the ant’s long-range visual receptor (Aksoy and Camlitepe, 2018) play a role in the context of controlling exploratory behaviours.

### A simple but functional heuristic

Oscillations in ants are optimized for visual exploration across azimuths (Clement et al., 2023) and consequently it would be useless to perform such exploratory behaviours in visually structure-less sceneries. Therefore, it makes functional sense that ants downregulate the production of oscillations in the presence of such structure-less environments (Fig. 2 G-H; Fig. 3B). The fact that ants rely on the presence of a diversity of edge orientations – rather than brightness or the actual quantity of edges – to control the production of oscillations (Fig. S2), can also be explained functionally. Indeed, relying on brightness or the overall number of edges within a visual scene would lead ants to oscillate and explore differently in visually cluttered environments (dark but large quantity of edges) and open environments (bright with little quantity of edges); even though both environments must be equally scanned across all azimuths to recall proper heading directions (Murray et al., 2020; Stürzl et al., 2016; Zeil et al., 2003). Relying on the presence of at least two different edge orientations appears as a good proxy to recognize an environment worth exploring. The only natural landscapes that would not fit in this criterion would be the ones with a perfectly flat horizon; in which case there would be only one type of edge orientation: the horizon. Such flat landscapes provide no directional information, so to repeatedly scan different directions is not beneficial. What’s more, ants tested in flat landscapes without terrestrial information spontaneously rely more on their path integration mechanisms to navigate to their goal in a straight line (Schultheiss et al., 2013; Schwarz and Cheng, 2010; Sommer and Wehner, 2004), that is, a navigation strategy that would be hindered, rather than helped, by the production of high amplitude lateral oscillations.

Perhaps more surprisingly, ants produced equally regular and prominent oscillations in a natural-like scenery than in an artificial scenery consisting of a same repeated pattern across azimuths, as long as the latter presented both horizontal and vertical edges (Fig. 3B). Such artificial repeated-pattern sceneries were directionally non-informative and thus are theoretically not worth exploring; yet the ants explored them thoroughly. This shows that ants pay no attention to whether the perceived scenery changes significantly across heading directions; or at least this information, which theoretically would be the best one to measure, is not used to control the production of oscillations. However, such environments, consisting of a perfectly repeated pattern of at least two types of edge orientation are extremely unlikely to exist in natural conditions. As it is often the case in ant navigation literature (Cheung et al., 2014; Cruse and Wehner, 2011; Menzel et al., 2005; Menzel and Muller, 1996; Wehner, 2003; Wehner and Menzel, 1990), we conclude that ants use a simple but efficient heuristic to decide whether a visual world is worth exploring. The latter seems to be a matched filter, enabling the ants to detect the presence of at least two different edge orientations.

We speak here of matched filter in the sense that ants are not ‘measuring’ the information that is directly related to their aim (i.e., oscillations are triggered accordingly to whether they currently provide pertinent visual information) but use a visual heuristic that matches natural conditions with the way they gather information through oscillations along the yaw axis (i.e., the presence of horizontal and vertical edges means oscillations should be useful given any natural environment). Surely, the match between the filter and the world is less impressive than in the context of sky compass orientation, where the receptive fields of neurons can literally match a natural distribution of the e-vector across azimuth and elevation (Homberg, 2004; Homberg et al., 2011). However, the visual terrestrial sceneries can vary in the natural habitat of *C. velox*, even over short distances: from arid landscape to meter high field of plants in spring and presence or absence of trees providing large skyline changes and even partial canopies above the ant’s nest (personal observations). Hence, assuming the presence of some horizontal and vertical edges may well fall close to the most optimized filter one could do to trigger exploratory behaviours.

How similar or different such a matched filter is in different ant species and whether it actually reflects some properties of their natural habitat, remains to be seen. North African *Cataglyphis fortis* ants living in saltpans usually forage in a featureless environment, relied more on their PI and were slower at learning visual cues as compared to *Melophorus bagoti*, which thrives in a more cluttered environment (Cheng et al., 2014; Schwarz and Cheng, 2010). Also, given that our experiments exclusively involved individuals raised in a lab environment, we cannot conclude whether the visual features identified here are developmentally constrained (Aepli et al., 1985; Bolzon et al., 2009; Ehmer and Gronenberg, 2002; Labhart and Meyer, 1999) and structural re-organization through pruning and synaptic change along visual pathways (Blakemore and Cooper, 1970; Cabirol et al., 2018; Grob et al., 2024; Gu and Kanai, 2014; Rössler, 2019; Rössler and Groh, 2012; Voss et al., 2017). Manipulating the visual environment during early experiences, as well as comparative studies across different ant species in such VR systems promises to shed light on the plasticity of the brain in interpreting a visual scenery.

### Neural considerations

Visual feature extractions can occur early in visual processing. Notably, lateral inhibition between neighbouring ommatidia extract local edges, which then can be integrated and summed up in the optic lobes (Bolzon et al., 2009; Srinivasan et al., 1982). From there, several pathways may convey various visual information to influence oscillations. Lateral oscillations in insects are likely produced in a pre-motor area called the Lateral Accessory Lobes (LAL), which receive multiple inputs from many brain regions (Chiang et al., 2011; Namiki et al., 2014; Namiki and Kanzaki, 2018, 2016a, 2016b; Steinbeck et al., 2020).

Regarding the effect of dynamic visual cues, wide-field rotational optic flow computed in the optic lobes (Busch et al., 2018; Egelhaaf, 2023; Pansopha et al., 2014) is locally compared to efference copies of the motor signal (Dauzere-Peres and Wystrach, 2023; Fenk et al., 2021; Kim et al., 2015), as well as proprioceptive feedback to form a prediction error signal that is sent to the LAL to modulate the oscillations (Dauzere-Peres and Wystrach, 2023). This control operates via an asymmetrical activation of the LAL enhancing a left or a right turn, in order to reduce the computed error. This is in congruence with the high amplitude turns (looping behaviour) observed in the absence of horizontal optic flow (Fig. 2G-L), which must result in a prediction error that activates one side of the LAL to prolong the current turn.

Our work reveals the existence of another pathway, which conveys this time information about static features in the visual scene, and notably, the simultaneous presence of at least two types of edge orientation in the scene. This pathway does not modulate the amplitude and frequency of oscillations but their actual presence and regularity in the outputted behaviour, which could be simply achieved through bilateral and thus overall excitation or inhibition of the LAL production of rhythmical oscillations. However, through which relay this pathway operates remains unclear. Evidence suggests the existence of a direct pathway from the optic lobe to the LAL (Namiki et al., 2014; Namiki and Kanzaki, 2018, 2016b; Steinbeck et al., 2020) providing a likely candidate to explain how the presence of a diversity of edges in the scene can modulate the activity of the LAL and thus the production of regular oscillations.

Alternatively, a parallel pathway exists from the optic lobe to the Mushroom Bodies (MBs) (Ehmer and Gronenberg, 2002; Gronenberg, 2001; Heisenberg, 2003; Paulk and Gronenberg, 2008), which outputs information to the LAL (Aso et al., 2014; Manjila et al., 2019; Scaplen et al., 2021; Steinbeck et al., 2020). The MBs are implicated in formation and retrieval of route memories of visual scenes in ants and can output a familiarity signal of the currently perceived scene (Buehlmann et al., 2020b; Kamhi et al., 2020; Webb and Wystrach, 2016; Wystrach, 2023), which we know can modulate the dynamics of oscillation (Clement et al., 2023; Murray et al., 2020). Whether information about the static visual feature identified here are conveyed through the MBs remains however *a priori* less likely, given that it operates already during early exploration of novel environments, without the need for learning.

Finally, the up- and down regulation of oscillations based on visual cues could also involve the Central Complex. This brain area keeps track of the insect current heading using mutli-modal information including visual and motion cues (Green et al., 2017; Kim et al., 2019; Stone et al., 2017; Turner-Evans and Jayaraman, 2016) and outputs left and right motor commands to the LAL to align the current heading with respect to a desired ‘goal heading’ (Beetz et al., 2023; Honkanen and Adden, 2019; Mussells Pires et al., 2024; Westeinde et al., 2024). We understand well how the Central Complex can adjust the course of ants in regards to both learnt (Matheson et al., 2022; Wystrach et al., 2020) and innate visual cues (Goulard et al., 2021) but how it is involved during the ant’s systematic search in an unfamiliar environment remains unclear. We could envision how the presence of visual cues such as the one characterised here could up- or down regulate the Central Complex influence on the ant’s motor control and hence modulate the intrinsic production of oscillations in the LAL.

## Conclusion

Here we show that ants adjust their visual exploration to the type of structure of the surrounding visual scenery. This control is based on a simple heuristic or ‘matched filter’ (Cheng and Freas, 2015; Wehner, 1987) detecting the simultaneous presence of both vertical and horizontal edges in the scenery (Grabowska et al., 2018), which upregulates the production of regular oscillations and thus exposes the ant’s gaze to multiple directions. Dynamic cues, such as optic flow, are also picked up but used for the different purpose of controlling the amplitude of the oscillations. How plastic and different this heuristic is across species remains unknown but the possibility to conduct experiments with navigating insects within VR systems promises to shed light onto how various species encode their visual natural environment, adapt to various early life experiences and thus helps to understand each species’ respective Umwelt (von Uexküll, 1934).

## Acknowledgements

We thank Hansjürgen Dahmen for providing us with the trackball system. We also thank Cody Freas, and Andrew Philippides for their helpful feedback on the manuscript. Funding: European Research Council, grant reference number: EMERG-ANT 759817, author: A.W.

## Supplemental information

**Figure S1:**
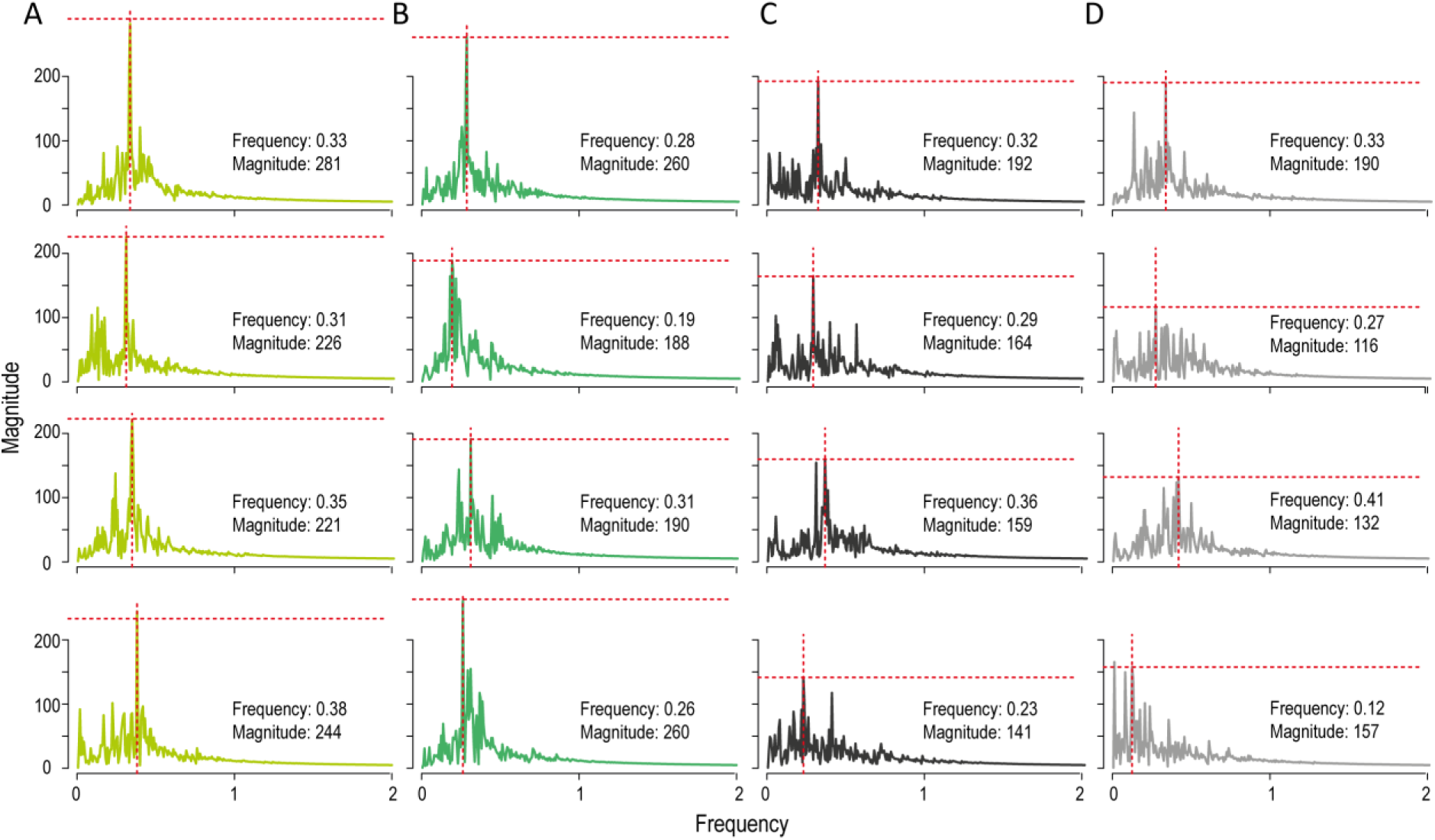
Power spectral density sample for different experimental manipulation. (A-D) Examples of ‘power spectral density’ four ants recorded with translational optic flow (TOF; A, light green) and without TOF (B, green) or recorded in visual surrounding made of vertical (C, grey) and horizontal (D, light grey) stripes. (A-D) Distribution of the individual Fourier dominant peak obtained from angular velocities times series across 160s. For each individual, the frequency peak with the highest magnitude was extracted, indicating a strong oscillation of the angular velocity signal at that frequency (dashed red lines).

